# Compression Sequencing enables ultra-sensitive and scalable scRNA-seq

**DOI:** 10.64898/2026.09.01.748706

**Authors:** Yan Yan, Mingjie Dai

## Abstract

Current sequencing methods are inefficient and bottlenecked by repeated sampling of highly abundant molecules, which dominate sequencing reads, limit assay throughput and sensitivity for rare targets. For example, single-cell RNA sequencing (scRNA-seq) can profile up to millions of cells, but remains severely constrained by sequencing cost, resulting in shallow gene coverage and high dropout rate. Here we report an information science-inspired method, “Compression Sequencing”, that tackles this fundamental inefficiency and enables highly improved (>100x) sequencing power. Our method works by performing an accurate and unbiased logarithmic transform on molecular abundances over a wide (5 logs) dynamic range, thus suppressing high-abundance targets and enriching rare ones, while maintaining quantitative accuracy. Applied to scRNA-seq libraries, our method allows ultra-sensitive detection of low-abundance transcripts (2-5x more UMIs), ultra-low sequencing cost (200x reduction), preserves accurate cell types and differential expression analysis over a 500-2,000 gene panel. In AML clinical samples, Compression Sequencing reproduces clinical diagnosis and additionally allows transcriptomic profiling at affordable cost (est. $10 per sample). Our approach thus enables ultra-sensitive and scalable single-cell analysis for large-scale functional genomics studies, drug discovery screens, AI cell model training, as well as affordable single-cell disease diagnostics.

Summary Fig.
Compression Sequencing enables ultra-sensitive and scalable scRNA-seq

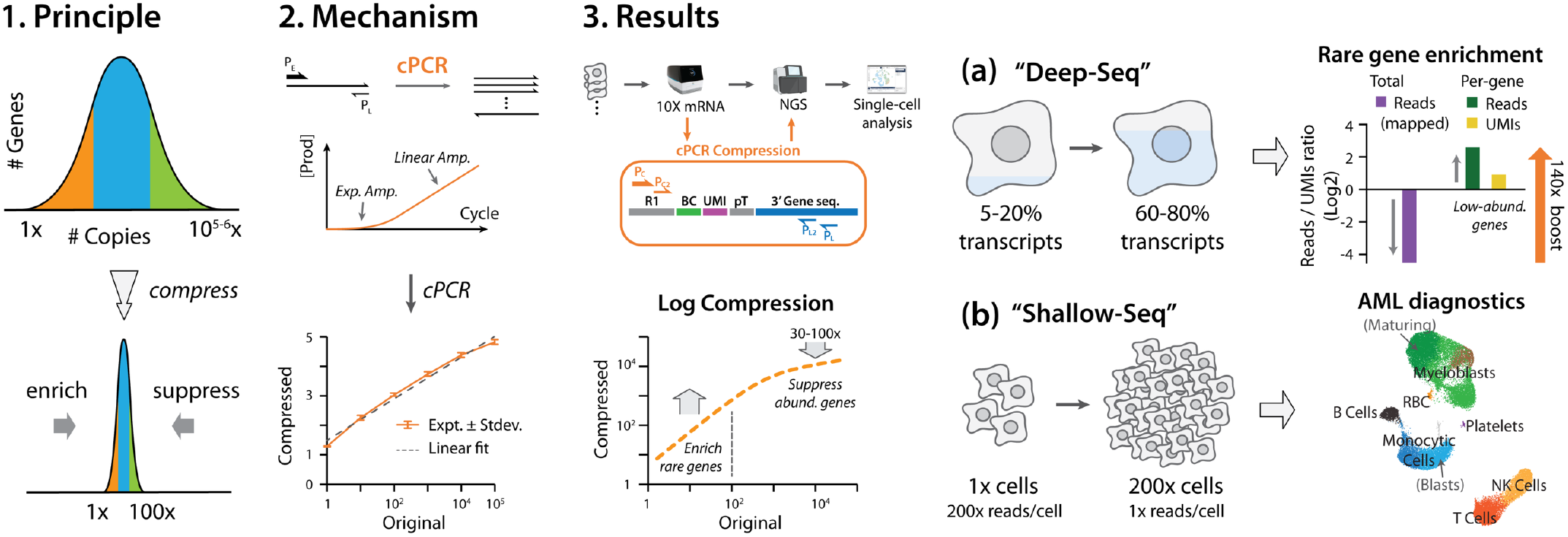

---

Single-cell RNA sequencing (scRNA-seq) methods have broadly transformed biological research and biomedicine^1–3^. Recent advances further enabled genome-wide functional studies^4–6^, tissue to organism cell atlases^7–9^, and large AI virtual cell models^10–14^. Whereas combinatorial and microfluidics-free cell barcoding techniques^15–17^ as well as decreasing sequencing cost^18^ allowed profiling of up to millions of cells^5,9,19^, current approaches sacrifice detection sensitivity and gene coverage for cell throughput^5,9,19^, posing significant limitations in detection sensitivity of rare genes and transcripts, as well as identification of small cell subpopulations^20–22^.

For example, many low-abundance transcripts exist in single-digit copies out of an estimated total of 300,000-500,000 mRNA molecules per cell^2,23,24^. Faithful detection of such rare transcripts requires a sequencing depth of 1-2 M reads per cell for full coverage^22^. However, current studies commonly allocate 10,000-20,000 reads per cell (50-100x lower), as a compromise between affordability, sensitivity and cell throughput^20,25,26^. Such shallow coverage translates to low gene sensitivity, high dropout rate, and an almost complete loss of single-cell heterogeneity in medium-to-low abundance genes (and their interaction patterns)^21^. On the other hand, functional genomic analysis, drug screens and virtual cell modeling require highly sensitive gene expression data, necessitating 100-500 cells per condition to achieve sufficient statistical power^4–6^. Additionally, it has been increasingly reported that, building a truly generalizable virtual cell model requires a training corpus under diverse cell lineage and perturbation conditions (at a scale of 30-100 M cells) with high-quality gene expression profiles^10-12,14^. However, even using the most advanced sequencers today, profiling 10 M cells at 50,000 reads/cell would take $120,000-250,000 for sequencing cost alone (UG100 or NovaSeq X).

Consequently, current single-cell sequencing methods are severely constrained by sequencing throughput and cost, reflected at both molecular and cellular levels, resulting in shallow gene coverage as well as limited sample throughput. Fundamentally, this is caused by the high dynamic range of mRNA expression in human cells^21,27,28^ (5-6 logs, Extended Data Fig. 1a). In a typical scRNA-seq study, genes below medium expression (70% of all genes) only take up a small fraction of all sequencing reads (<1%), and shows high (50-100%) dropout rates^23,29^ (Extended Data Fig. 1b). On the other hand, a small fraction of highly expressed genes (<10%) take up most (>95%) of sequencing reads, which contribute little to cell type determination or gene expression analysis, but constrain the required sequencing depth per cell and total cell throughput.

Several approaches for tackling the sequencing throughput limitation have been previously reported, such as using targeted gene panel enrichment^30–32^, hybridization kinetics-based normalization^33–35^ or duplex-specific nucleases (DSN) digestion^36,37^. Unfortunately, these approaches either provided insufficient reduction in dynamic range, or are prone to sequence-dependent biases, resulting in limited improvement of sequencing efficiency or compromised quantitation accuracy.

Here we report a biochemical solution to this problem, that applies a dynamic-range-compressing transformation to molecular abundances while preserving quantitative readout (Fig. 1a-c, Extended Data Fig. 2a). Similar transform has been foundational in computer science and digital signal processing, enabling efficient data storage^38–40^ and signal transmission^41^ (Fig. 1c). Applied to complex sequences libraries such as in scRNA-seq, such an transform will redistribute sequencing reads from highly abundant genes to rare ones, thus improving sequencing efficiency and sensitivity, and allowing accurate gene expression analysis across a wide range of expression levels (Fig. 1d-e).

**Figure 1.**
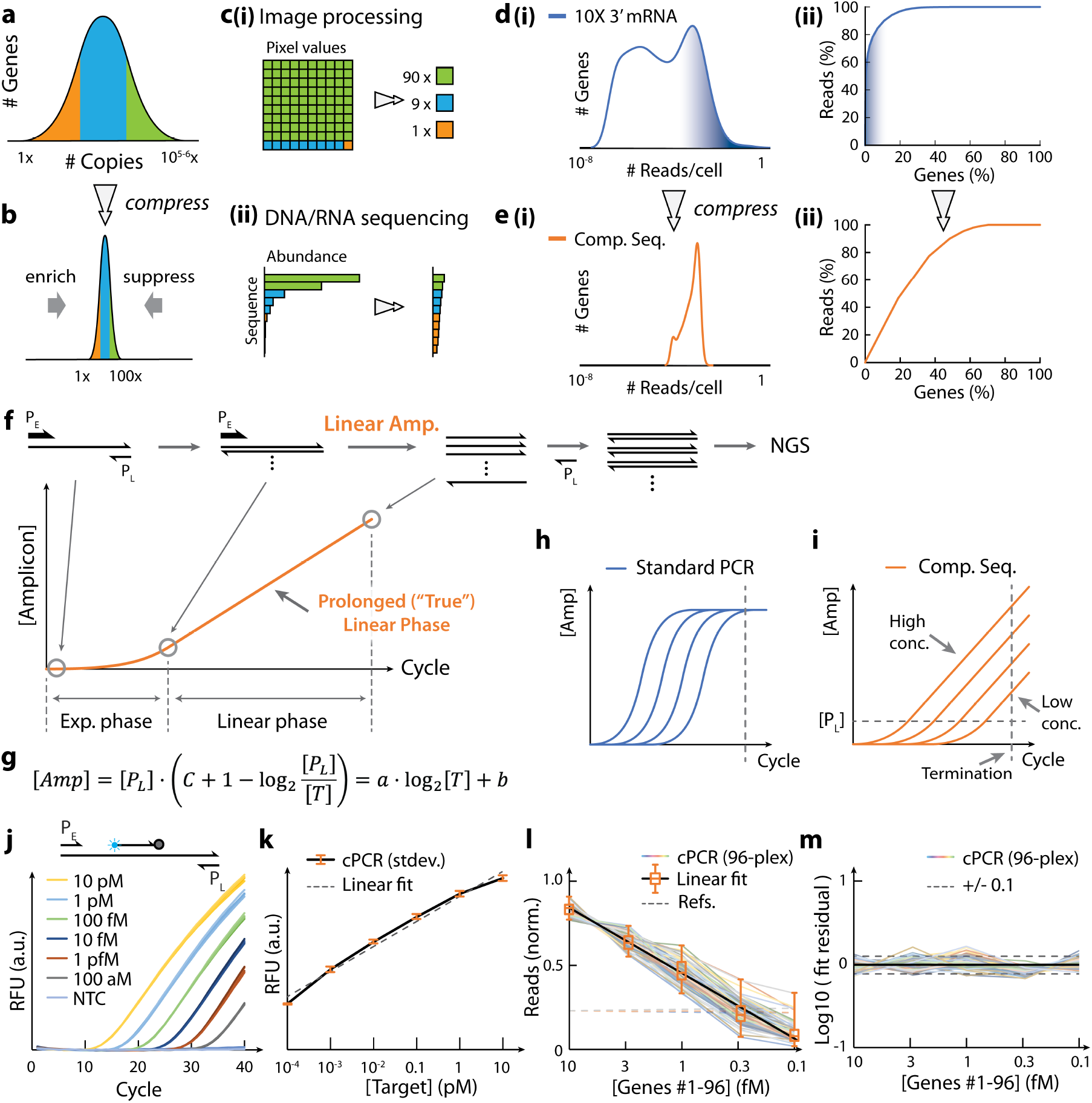
Compression Sequencing enables accurate logarithmic transform and dynamic range compression in highly multiplexed assay. (**a-b**) Principle of dynamic range compression in complex sequence library, showing gene abundance distribution in (**a**) original and (**b**) compressed library. High-abundance targets are suppressed; rare targets are enriched. (**c**) Same principle of data compression applies to (**i**) digital image processing and (**ii**) high-throughput sequencing. (**d**) (**i**) Gene expression distribution (shown as kernel plot) in a representative scRNA-seq dataset (10X 3’ mRNA, in human PBMC cells). (**ii**) Resulting reads allocation among all genes. Shaded areas represent a small fraction of high-abundance genes that dominate sequencing reads, and constrains sequencing efficiency, sensitivity and cost. (**e**) Gene expression distribution (**i**) and reads allocation (**ii**) using Compression Sequencing (simulated). Compression Sequencing allows near-uniform allocation of reads across genes of different abundance levels. (**f**) Principle of Compression PCR (cPCR). cPCR uses different concentrations for forward and reverse primers, and proceeds in two phases: an initial exponential phase followed by a (prolonged) linear phase. Single-stranded cPCR product is converted to dsDNA before library prep and sequencing. (**g**) cPCR conversion formula. [P_L_], [T], [Amp] denote concentrations of the limiting primer, input target sequence, and cPCR product (sum of dsDNA and ssDNA), respectively; C denotes total number of PCR cycles; a and b are constant parameters (Supplementary Note 1). (**h-i**) Comparison of conventional PCR (**h**) and cPCR (**i**) amplification traces at different input target levels. cPCR allows uniform and accurate quantitation across a wide dynamic range. Proof-of-concept cPCR test. Plot shows real-time cPCR amplification traces monitored by qPCR assay, confirming a two-phase reaction. Three replicates are shown for each input level. (**k**) End-point qPCR signal as a function of input target level shows accurate log transform over a wide dynamic range (5 logs); dashed line indicates best linear fit. (**l-m**) 96-target multiplexed Compression Sequencing shows highly accurate log transform under complex genomic background. (**l**) Sequencing reads vs target concentration (R^2^ > 0.95). (**m**) Fitting residual vs target concentration (<30% CV after conversion to linear scale). In (l) and (m), each colored line represents one of the 96 targets. In (l), box center and edges represent median and quartiles, whiskers represent most extreme data points; dashed lines represent spiked-in references. In (m), dashed lines represent +/-0.1 offset as visual guides. See Methods for analysis details.

Our method, “Compression Sequencing”^*^, employs a multiplex PCR-based reaction, which performs an accurate logarithmic transform of sequence abundance over a wide range (5 logs) of input (up to 10,000x compression). Applied to PBMC scRNA-seq libraries, Compression Sequencing allows accurate and unbiased sequence compression over hundreds to thousands of genes in parallel, achieving ultra-sensitive detection (100x sensitivity, 2-5x UMIs in rare genes) and scalable single-cell profiling (200x lower cost). Additionally, Compression Sequencing correctly identifies disease cell states and reports subtype diagnosis in AML clinical samples, at up to 500x reduced sequencing depth and cost ($<10 per sample), suggesting affordable scRNA-seq based diagnostics with transcriptomic analysis. Our method is compatible with multiple commercial single-cell workflows (e.g. 10X Genomics, Fluent/Illumina), and has the potential to enable broad applications from basic research to clinical settings, e.g. large-scale genetic and drug screens, organism and development cell atlas, AI virtual cell and disease models, and single-cell disease diagnostics.

## Results

### Compression PCR is an accurate molecular logarithmic encoder

Here we report a novel, multiplex PCR-based approach, “Compression PCR (cPCR)”, that performs dynamic range compression on molecular abundances and functions as a “molecular logarithmic encoder”. Specifically, our method builds on the concept of asymmetric PCR^42,43^, where the forward and reverse primers are supplied in different concentrations in a PCR reaction, but with several critical design adaptations (Fig. 1f-i). Under specially designed experimental conditions, instead of a conventional exponential growth (single-phase), the reaction proceeds through a two-phase amplification process: (i) an initial, exponential phase, where conventional PCR takes place with both primers available, followed by (ii) a second, linear phase, where the “limiting primer” (P_L_) has been depleted, and the reaction continues to produce ssDNA using the “excess primer” (P_E_) only (Fig. 1f). Here, the second phase (“prolonged linear phase”) reflects true linear growth due to single-primer amplification, which is different from the transient “linear phase” observed in conventional qPCR assays.

At the end of the reaction, the amplicon concentration (predominantly ssDNA) can be determined by a logarithmic transform of the initial template concentration (Fig. 1g, Supplementary Note 1), thus achieving an accurate, molecular logarithm conversion. When applied to thousands of genes in a complex sequence library, this reaction further allows unbiased compression in sequence abundances, preferentially enriching rare sequences while suppressing high-abundance ones (Extended Data Fig. 2a). Compared to conventional PCR (either with early or late reaction termination), cPCR uniquely allows dynamic range compression and quantitation over a wide range of input concentrations (Fig. 1h-i, Extended Data Fig. 2b-c).

Critically, our cPCR method has three unique advantages:

i. (1)_ cPCR is unbiased, i.e. the compression mechanism works the same way for low- or high-abundance genes, without relying on any presumption of their expression levels.
ii. cPCR is quantitative and independent for each gene, i.e. cPCR performs logarithmic compression independently in a multiplexed reaction, and preserves quantitative expression for each gene.
iii. cPCR is tunable, i.e. the degree of dynamic range compression can be controlled by adjusting P_L_ concentration and other reaction conditions, to achieve an optimal balance between sequencing cost reduction and detection sensitivity improvement for rare targets.

We first performed *in silico* analysis on the performance of dynamic range compression and sequencing efficiency improvement using our method. Our analysis (Extended Data Fig. 1a-c) showed that, on a representative scRNA-seq dataset, logarithmic compression suppresses read fraction from high-abundance genes by 40x while boosting that from low-abundance ones, achieving up to 500x overall improvement in sequencing efficiency and sensitivity in rare genes (Extended Data Fig. 1g,h). Similar improvement can also be observed in panel-based assays using targeted enrichment (Extended Data Fig. 1d-f).

We performed a proof-of-principle cPCR test on synthetic amplicons to assay the experimental performance in dynamic range compression (Fig. 1j). Using an excess primer ratio (P_E_/P_L_) of 50, and a TaqMan probe readout (against the P_E_ strand), we observed a clear two-phase amplification process, over 5 logs of input concentration. Our results showed a logarithmic transform from 5 logs to <1 log (>10,000-fold reduction), along with a high signal accuracy across the entire range of input levels (R^2^ = 0.99), allowing accurate and unbiased compression (Fig. 1k). Next, we tested multiplexed cPCR under a four-target setting, and varied the concentration of two targets from 10 pM to 10^-4^ pM while keeping the other two fixed. We found that, Taq polymerase combined with inhibitor-tolerant reaction buffer (Bio-Rad SsoAdvanced) provide the cleanest compression for multiplexed cPCR. Under optimized buffer and reaction conditions, our results showed independent and accurate logarithmic compression for all four targets (Supplementary Fig. 1a-b), with high signal linearity (R^2^ = 0.98-0.99) and little interference across targets (Supplementary Fig. 1c).

### Compression Sequencing allows highly multiplexed, unbiased compression in complex sample

We developed a Compression Sequencing workflow based on the cPCR reaction, by converting predominantly single-stranded cPCR product to double-stranded form with a primer extension reaction using P_L_ (Fig. 1f), followed by ligation-based library prep and sample indexing. Additionally, reference sequences can be spiked in at the double-strand conversion step, for concentration calibration and sample normalization.

We first validated the workflow using an 8-target multiplexed test with lowered limiting primer concentration (15 nM) and primer ratio (P_E_/P_L_ = 10), to be compatible with highly multiplexed tests later. We noticed that insufficient polymerase and reagent usage results in incomplete reaction, whereas excessive supply can induce off-target amplification. After systematic tests, we established a generalized biophysical model for polymerase usage based on primer concentration, target multiplexity and annealing time (Supplementary Fig. 2, Supplementary Note 2). Under optimal conditions, our results showed highly accurate compression in all targets (R^2^=0.94-0.99) over a high dynamic range (5 logs, Supplementary Fig. 2a). Moreover, our model predicts optimal reagent usage over a range of different cPCR reactions (up to 16-target tests), maintaining highly accurate compression in all tested cases (Supplementary Fig. 2g-i).

To further scale up Compression Sequencing for highly multiplexed compression over hundreds to thousands of sequences, we designed larger primer pools that are simultaneously optimized in (i) primer binding thermodynamics^44^, taking into consideration of P_E_/P_L_ imbalance, and (ii) minimal primer dimer formation, adapted from a Simulated Annealing algorithm^45^ (Extended Data Fig. 2d). Using stringent design criteria and higher iterations, our design achieved >10x reduced primer dimer formation compared to previous report^45^ (Extended Data Fig. 2e).

We tested multiplexed cPCR on a panel of 96 energy-balanced primers against human gene targets under genomic background. Without individual primer optimization, 69 of 96 targets (72%) showed accurate logarithmic transform (R^2^>0.95) across the entire tested concentration range from 10 fM to 100 aM; 83% targets showed R^2^>0.90 (Fig. 1l). Additionally, cPCR showed accurate relative abundance (CV<30%, normalized by peak signal, Fig. 1m). We further performed a two-species mixture test on 96 human and 96 yeast genes (192 total targets), with one group (human) kept at a fixed concentration and the other (yeast) varied from 10 fM to 100 aM (Extended Data Fig. 2f). Our results showed that, cPCR maintained constant signal level for the fixed group, while achieving accurate logarithmic signal compression on the varied group (R^2^>0.98, Extended Data Fig. 2g).

### 1,000-fold dynamic range compression and accurate cell type-specific analysis in scRNA-seq library

Compression Sequencing promises to greatly enhance scRNA-seq efficiency by reducing the wide dynamic range in gene expression (5-6 logs, Fig. 1d). We developed an “add-on” protocol for applying Compression Sequencing to scRNA-seq libraries (3’ capture, Fig. 2a-b). We placed the gene-specific limiting primer (P_L_) against the mRNA transcript, 100-200 nt upstream from the poly-A site; and designed a common excess primer (P_C_) within the Illumina read primer on the opposite side, so that the amplicon includes the cell barcode, UMI and a segment of gene-specific sequence (Fig. 2a). To facilitate scaling up to hundreds of gene targets and maintain accurate cPCR compression in complex transcriptomic background, we made further developed a nested cPCR strategy by replacing double-stranded conversion step with a second (regular) PCR, using a pair of nested primers (P_L2_ P_C2_) (Fig. 1f, Fig. 2a). Finally, since the common primer is shared among all genes and will be supplied in significant excess (e.g. for 500 genes, P_C_/P_L_ ratio = 500×15 = 7,500), we carefully modelled primer binding thermodynamics and minimized dimer formation tendency using stringent parameters during primer design.

We tested Compression Sequencing on a scRNA-seq library (10X 3’ mRNA) of a human peripheral blood mononuclear cell (PBMC) sample.

**Figure 2.**
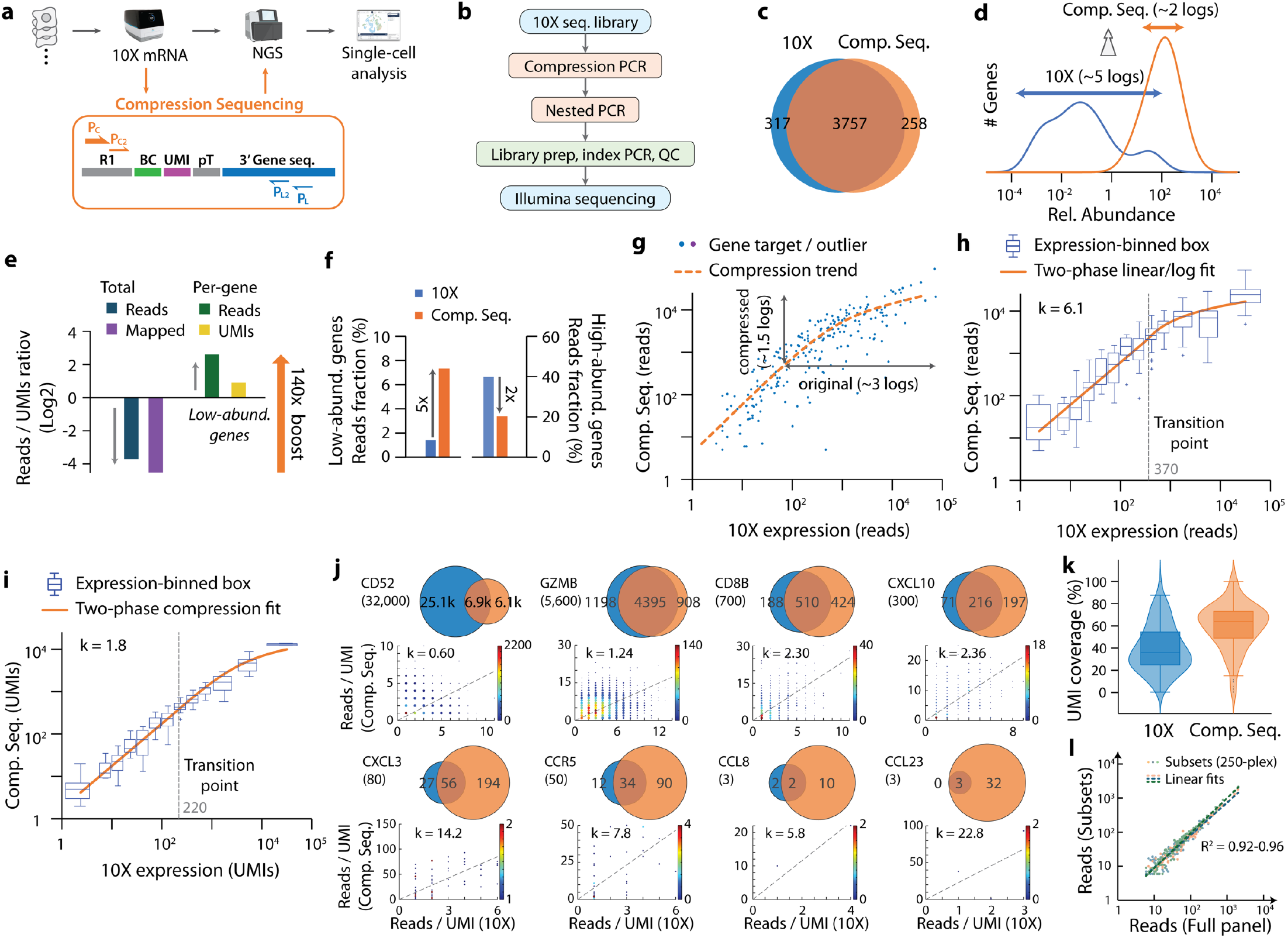
Compressed “Deep Sequencing” with >100x improved efficiency reveals 2-5x more transcripts in low-abundance genes. (**a-b**) Schematic workflow (**a**) and key steps (**b**) of Compression Sequencing adapted to 10X Genomics single-cell mRNA expression assay. Outer primers (P_C_ and P_L_) are used for cPCR reaction, inner primers (P_C2_ and P_L2_) are used for nested PCR. (**c-d**) Compression Sequencing shows near complete cell barcode coverage (**c**), and an overall ~1,000x dynamic range compression (**d**). (**e-f**) Effect of sequence compression in (**e**) total reads reduction and overall enrichment, and (**f**) reads re-allocation between low-abundance and high-abundance gene fractions. In (e), enrichment in low-abundance gene targets shown for reads and UMIs, respectively. All ratios are shown in log2 scale, with a 140x overall enrichment. (**g**) Reads compression plot, comparing Compression Sequencing vs 10X reads for each gene, with visual guides showing two-segment reads conversion: linear amplification for low-abundance genes followed by logarithmic compression for high-abundance targets. (**h**) Same reads plot as in (g), shown as abundance-binned box plots after outlier removal, along with fit (solid line) to a two-segment reads compression model (Supplementary Note 3). (**i**) UMI conversion plot, binned and shown as box plots similarly as in (h), along with fit (solid line) to a two-segment UMI conversion model: linear amplification followed by reads-based stochastic sampling (Supplementary Note 3). In (h) and (i), box center and edges represent median and quartiles, whiskers extend to most extreme data points not including outliers. (**j**) UMI coverage analysis in Compression Sequencing, showing 8 examples in genes with different abundance levels. For each gene, top-left: gene name and abundance (UMI count in 10X dataset); top-right: UMI overlap between 10X (blue) and Compression Sequencing (orange); bottom: UMI-resolved reads comparison shown as dot-based heat map (size and color represents UMI degeneracy), dashed lines indicate best linear fit through origin. (**k**) Estimated UMI coverage (%) using Compression Sequencing vs 10X, estimated from UMI overlap between the two datasets. (**l**) High accuracy of Compression Sequencing, comparing full gene panel (500 genes) vs three different subsets (250 genes each). Dotted lines represent linear fits. See Methods for analysis and fitting details.

As the reference dataset, we sequenced the 10X library to 33,000 reads/cell (140 M reads, 4,200 recovered cells), then performed standard analysis routines for clustering and cell type annotation (Supplementary Fig. 3a, Supplementary Fig. 4a-d). On a pilot set of 60 gene targets (without nested PCR), Compression Sequencing showed near complete cell barcode coverage (>99% from 100,000 reads, Fig. 2c), as well as 1,000-fold reduction in gene expression dynamic range (Fig. 2d).

We next designed a larger test panel consisting of 148 targets covering the full dynamic range (5.5 logs). Our results showed that, nested cPCR significantly boosted on-target read fraction (from 40% to 70%), and produced a clear logarithmic transform across the entire dynamic range (Supplementary Fig. 3b), showing 500-1,000x suppression in high-abundance genes and 5-10x enrichment in rare ones. Additionally, Compression Sequencing showed highly reproducible signal between technical replicates (R^2^=0.99, Supplementary Fig. 3c).

After single cell clustering, we mapped the cell barcodes onto the same UMAP projection as in the reference dataset, taking advantage of the “add-on” workflow. Our results showed that, Compression Sequencing preserves specific marker gene expression, accurate cell type annotation (Supplementary Fig. 3d), and reproduces differential expression analysis (R^2^=0.94, Supplementary Fig. 3e). Moreover, Compression Sequencing reports more UMIs and higher cell barcode coverage in medium to low abundance genes (Supplementary Fig. 3f).

### Compressed “Deep Sequencing” with >100x improved efficiency reveals 2-5x more transcripts in low-abundance genes

To quantitatively assay the performance of Compression Sequencing in highly multiplexed assay, we designed a 500-gene immune signature panel. We expanded the panel to cover unique 3’-end sequences from all annotated splicing isoforms in RefSeq database (486 genes, 533 unique 3’-end sequences), which produced a set of 512 cPCR primer pair sequences (success rate 93%, covering 457 genes). We further lowered P_L_ concentration to 7.5 nM to accommodate for highly multiplexed reaction, and extended cPCR annealing time to balance hybridization kinetics.

We sequenced the compressed library to a depth of 11 M reads (~1/13x of 10X reference dataset, Fig. 2e, Supplementary Fig. 4e-h), which we termed “Deep Sequencing” relative to “Shallow Sequencing” later. At similar saturation levels, our results showed greatly improved sequencing efficiency (100-140x), e.g. reduced from 200 M to 2 M at 0.60 saturation (Extended Data Fig. 3a). Extrapolation using the preseq software^46^ up to 10^12^ total reads also suggested a 100x reduction in library complexity (Extended Data Fig. 3b-c), consistent with the observation above. Read fraction analysis on subsets of high-quality gene targets (Fig. 2f) further showed that, library compression boosts read fraction of low-abundance genes (from 1.5% to 7.4%, 5x increase), and suppresses that of high-abundance targets (from 43% to 22%).

We next performed quantitative analysis on library compression accuracy and detection sensitivity in individual genes. Scatter and binned read plots against 10X reference dataset (200 nt capture range, Fig. 2g-h) showed a clear two-segment compression behavior: linear read amplification for low-abundance targets (which did not finish exponential phase during cPCR), and logarithmic compression for high-abundance ones (which successfully transitioned into linear phase). Fitting to a two-segment mathematical model (Supplementary Note 3) suggested a transition point at 370 reads (in 10X dataset), equivalent to 150 UMIs (0.035/cell). Notably, cPCR performed 30x reduction of dynamic range in mid-to-high expression genes, at the same time reallocated the saved reads to maximally preserves detection sensitivity in rare genes (no compression, 6x reads boost, achieving an overall 140x enrichment, Fig. 2e).

Deeper reads allows more transcript (UMI) detection. We investigated UMI detection as a function of gene abundance, using a similar two-segment model as above, but with an additional stochastic sampling step (Fig. 2i, Supplementary Note 3). Our results showed an average of 1.9x boost in detected UMIs (up to 5x) in low-abundance genes (Fig. 2e), consistent with an estimated 4x UMIs from preseq modelling (Extended Data Fig. 3b). Notably, even in medium-abundance genes, Compression Sequencing maintained a high UMI coverage despite significant reads compression, facilitating accurate downstream expression analysis. We further performed UMI-resolved reads enrichment analysis in medium and low abundance genes. Our results showed an unbiased, independent read conversion for each gene target: lower enrichment for high-abundance genes; higher amplification (20-50x) along with deeper UMI detection (3-10x) in low-abundance targets (Fig. 2j). Overall, Compression Sequencing achieves consistently higher UMI coverage (median 64%) compared to 10X reference dataset (median 36%) (Fig. 2k).

Finally, we investigated signal stability and technical reproducibility of our method. We observed highly reproducible compressed reads between technical replicates (R^2^ > 0.98, Extended Data Fig. 3e), as well as between full panel and half-sized subsets (R^2^ = 0.92-0.96, Fig. 2l). Additionally, we observed high consistency in compressed reads using primer set designs (R^2^ = 0.85-0.90, Extended Data Fig. 3f). Combined, these results validate the high accuracy and robustness of Compression Sequencing in a highly multiplexed scRNA-seq setting.

### Comprehensive immune profiling in a 2,000-gene panel allows more sensitive and reproducible cell type-specific DEG detection

We next applied Compression Sequencing for comprehensive immune profiling on a 2,000-gene panel. The panel consists of a total of 2,086 immunology markers and signature genes (18.7% of total abundance, 2,395 unique 3’ sequences), split into four parallel cPCR sub-panel of ~500 genes each, with 186 repeats as cross-panel validation (Fig. 3, Extended Data Fig. 3g). Compressed “Deep Sequencing” (51 M reads, ~1/3x of 10X reference) showed expected reduction in library complexity (Extended Data Fig. 3g) and reads fraction re-distribution from high-abundance genes (4.4x reduction) to the low-abundance side (5.1x enrichment, Extended Data Fig. 3h-i), similar to the 500-gene experiment above.

**Figure 3.**
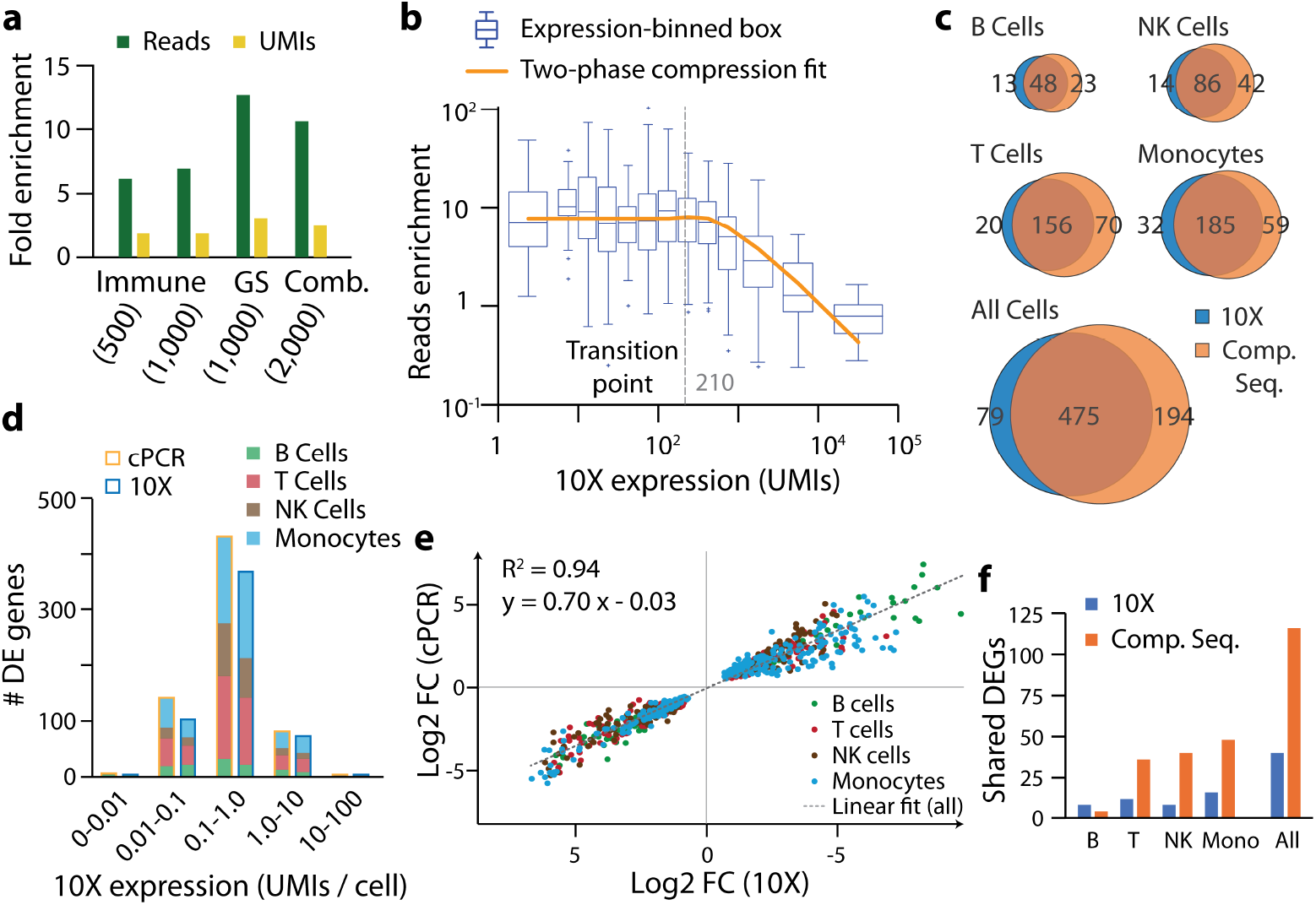
Comprehensive immune profiling in a 2,000-gene panel allows ultra-sensitive and reproducible DEG analysis. **(a)**Reads and UMIs enrichment in low-abundance genes, using Compression Sequencing, shown for different gene panels and sub-panels: Immune 1,000-gene panel and 500-gene sub-panel, Gene signature 1,000-gene panel, and combined 2,000-gene panel. (**b**) Reads enrichment ratio shown as abundance-binned box plots, along with fit (solid line) to a two-segment model: constant amplification for low-abundance genes and compression for high-abundance targets (Supplementary Note 3). Box center and edges represent median and quartiles, whiskers extend to extreme data points not including outliers. (**c**) Differentially expressed genes (DEGs) detection overlap between Compression Sequencing and 10X, shown in four major cell types and in sum. (**d**) DEG coverage broken down by cell type (face color) and gene abundance bins (x axis), showing higher boost in DEG detection in lower abundance bins, using Compression Sequencing. (**e**) Log2 fold change (Log2FC) plot between Compression Sequencing and 10X assay, in overlapping DEGs. Colors indicate cell types; dashed line indicates best linear fit (equation shown in figure). (**f**) Comparison of shared DEGs (detected in two independent samples) in four major cell types, showing more sensitive and reproducible DEG detection using Compression Sequencing. See Methods for analysis and fitting details.

Compression analysis using the two-segment model showed highly sensitive detection in rare genes (10.6x reads boost, 2.5x UMIs, Fig. 3a, Extended Data Fig. 3j-k) compared to the reference dataset, achieving an overall 30x reduction in dynamic range (Fig. 3b). UMI coverage analysis further confirmed significantly higher transcript detection sensitivity (67% vs 27%) (Extended Data Fig. 3l). Moreover, cross-panel comparison showed highly reproducible gene detection (Extended Data Fig. 3m) with minimal off-panel cross-talk (Extended Data Fig. 3n-o).

Depper UMI coverage allows more sensitive and robust differential expression (DE) analysis. In the PBMC sample, our results showed 21% higher total DEG detection in four major cell types (B, T, NK and Monocytes, 12-28% per cell type) within the gene panel (Fig. 3c). Notably, the extra DEGs exhibit a clear enrichment in medium-to-rare genes that are difficult to detect reliably (<1.0 UMIs/cell, Fig. 3d), including low-expression transcription factors, cytokine/chemokine, surface receptors and signaling proteins. Among overlapping DEGs, our results showed highly consistent analysis with 10X reference (Log2FC R^2^=0.98, lower slope likely due to ratio inflation from close-to-zero UMI counts^47^ in 10X dataset, Fig. 3e, Extended Data Fig. 3p)

We further asked if Compression Sequencing allows reproducible DEG analysis across biologically independent samples. We performed similar paired standard and compressed scRNA-seq, following the same add-on workflow (10X 3’ mRNA, then Compression Sequencing, Fig. 2a) using a different donor sample (Sample 2, Extended Data Fig. 4). Compression Sequencing faithfully reproduced the logarithmic reads compression (Extended Data Fig. 4a-c), along with 10x reads enrichment and 3x UMI coverage in low-abundance genes (Extended Data Fig. 4d-f).

DEG analysis among four major cell types reported 30% higher total detection (up to 45% per cell type, Extended Data Fig. 4g-h), and high agreement with 10X dataset in overlapping DEGs (Extended Data Fig. 4i-j); additionally, showing an excellent agreement across two donor samples (Log2FC R^2^=0.96, slope=1.00, Extended Data Fig. 5b). Furthermore, Compression Sequencing showed considerably higher cross-sample reproducibility (Extended Data Fig. 5c): we observed 45% overlap in DEGs reported only by Compression sequencing, in contrast to 21% in 10X-only genes; in total we observed 3x higher DEG count (Fig. 3f). Our results showed that, by compressing library complexity and improving UMI coverage, Compression Sequencing allows more sensitive and robust DEG analysis, even with considerably reduced sequencing depth.

### Compressed “Shallow sequencing” allows scalable scRNA-seq with 200x reduced depth

We next tested if the high sequencing efficiency provided by sequence compression can be repurposed to enable cost-effective scRNA-seq with significantly reduced depth. We performed “Shallow Sequencing” on the compressed scRNA-seq library (500-gene immune panel) at a depth of 0.7 M reads (200x reduced from 10X dataset). After unsupervised clustering and marker-based cell type assignment, we observed near complete (>96%) cell barcode coverage and highly accurate cell type identification (Fig. 4a, mapped to 10X projection), with an adjusted random index (ARI) of 0.78 among the four major cell types (B, T, NK and monocytes; ARI=0.97 if combining T and NK cells).

**Figure 4.**
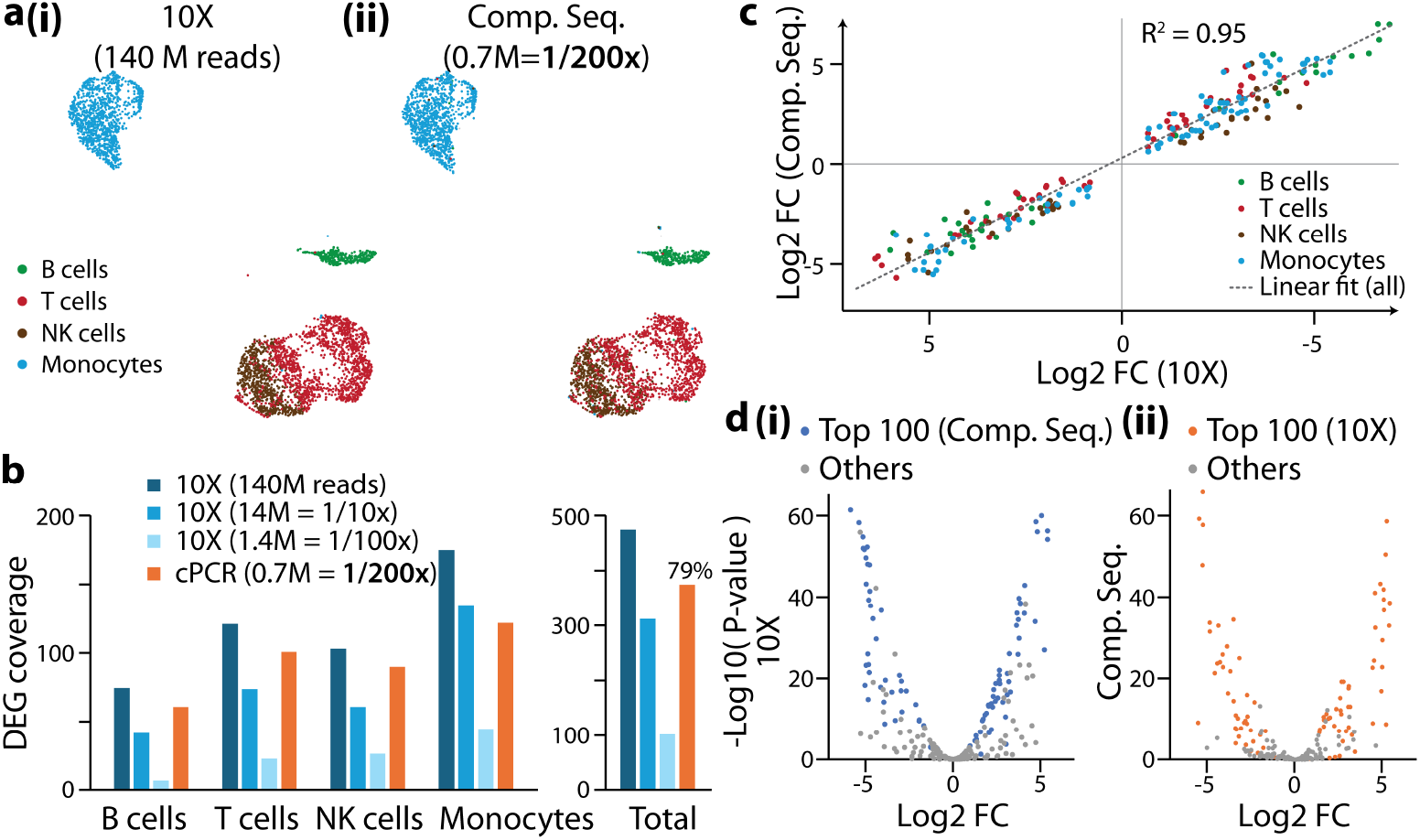
Compressed “Shallow Sequencing” with 200x lower depth enables ultra-scalable scRNA-seq. (**a-d**) Compressed Shallow Sequencing using 1/200x reads compared to 10X dataset (0.7 M vs 140 M) in PBMC scRNA-seq library, using a 500-gene immunology panel. (**a**) UMAP projection comparison of cell clustering and cell type assignment in (**i**) 10X and (**ii**) Compression Sequencing datasets. Compression Sequencing dataset is mapped to 10X projection; same colors represent same cell types in both maps. (**b**) Comparison of DEG coverage in 10X dataset (original, 1/10x and 1/100x subsampled) and Compression Sequencing (at 1/200x depth), in four major cell types (left) and combined (right). (**c**) Differential expression (Log2FC) comparison between Compression Sequencing and 10X dataset, in overlapping DEGs. Colors indicate cell types; dashed line represents best linear fit (R^2^=0.95, slope=0.95). (**d**) Volcano plots for differential expression analysis in monocytes in (**i**) 10X and (**ii**) Compression Sequencing datasets. In both plots, colored dots (blue or orange) represent 100 most significant genes (lowest P-value) detected by the other method; gray dots represent all other genes. See Methods for analysis details.

Differential expression analysis using Compression Sequencing showed a high coverage (79%) of in-panel DEGs reported by the reference dataset, significantly outperforming sub-sampled 10X datasets at equivalent depth (<15% at 1/100x depth). Additionally, among overlapping DEGs, Compression Sequencing reported highly accurate differential expression in all cell types (Log2FC R^2^ = 0.95, slope = 0.95, Fig. 4c), and showed high agreement with 10X in high-confidence DEG detections (Fig. 4d).

Combined, our results suggest that efficient sequence compression allows faithful scRNA-seq at significantly reduced sequencing reads and cost (1/200x from 10X, <170 reads/cell), while preserving quantitative gene expression information, allowing accurate cell type identification as well as differential expression analysis.

### Affordable scRNA-seq testing for AML subtype diagnostics with transcriptomic profiling

Acute Myeloid Leukemia (AML) is an aggressive hematologic malignancy with a poor 5-year overall survival rate of ~30%^48,49^. Recent single-cell sequencing studies have revealed previously unrecognized cellular sub-populations and disease subtypes^50–53^, and enabled earlier detection of measurable residual disease (MRD)^54^. Despite these advances, high cost and limited scalability of current scRNA-seq assays remain major barriers to its clinical deployment.

Compression Sequencing shows promise for affordable single-cell testing in diseases such as AML. To explore this possibility, we applied our method to three clinical PBMC samples associated with different AML subtypes and treatment history (S1-S3, two paired M4 and one M2 subtypes, Extended Data Fig. 6a), along with a healthy donor control (S0) (Fig. 5a). We performed single-cell barcoding using Fluent (Illumina) PIP-seq kit^17^, to allow scalable profiling with affordable cell barcoding cost. We employed a similar add-on workflow (as in Fig. 2a) and performed standard sequencing (38,000 reads/cell, as the “reference” datasets) for the unmodified PIP-seq library, followed by Compression Sequencing and shallow sequencing. Sequencing data were first converted to 10X style, then aligned and further analyzed using Cellranger and custom program. After co-clustering the reference datasets (S0-S3, prepared with Fluent) with two healthy donor samples (prepared with 10X) using Harmony, we observed a clear three-segment differentiation landscape showing myeloblast, monocytic and lymphoid compartments (Extended Data Fig. 6b). All three healthy donor samples (two Fluent, one 10X) showed a high degree of overlap in all cell types. In contrast, AML disease samples (S1-S3) exhibited abundant subpopulations of blasts outside of healthy cell clusters (Extended Data Fig. 6c).

**Figure 5.**
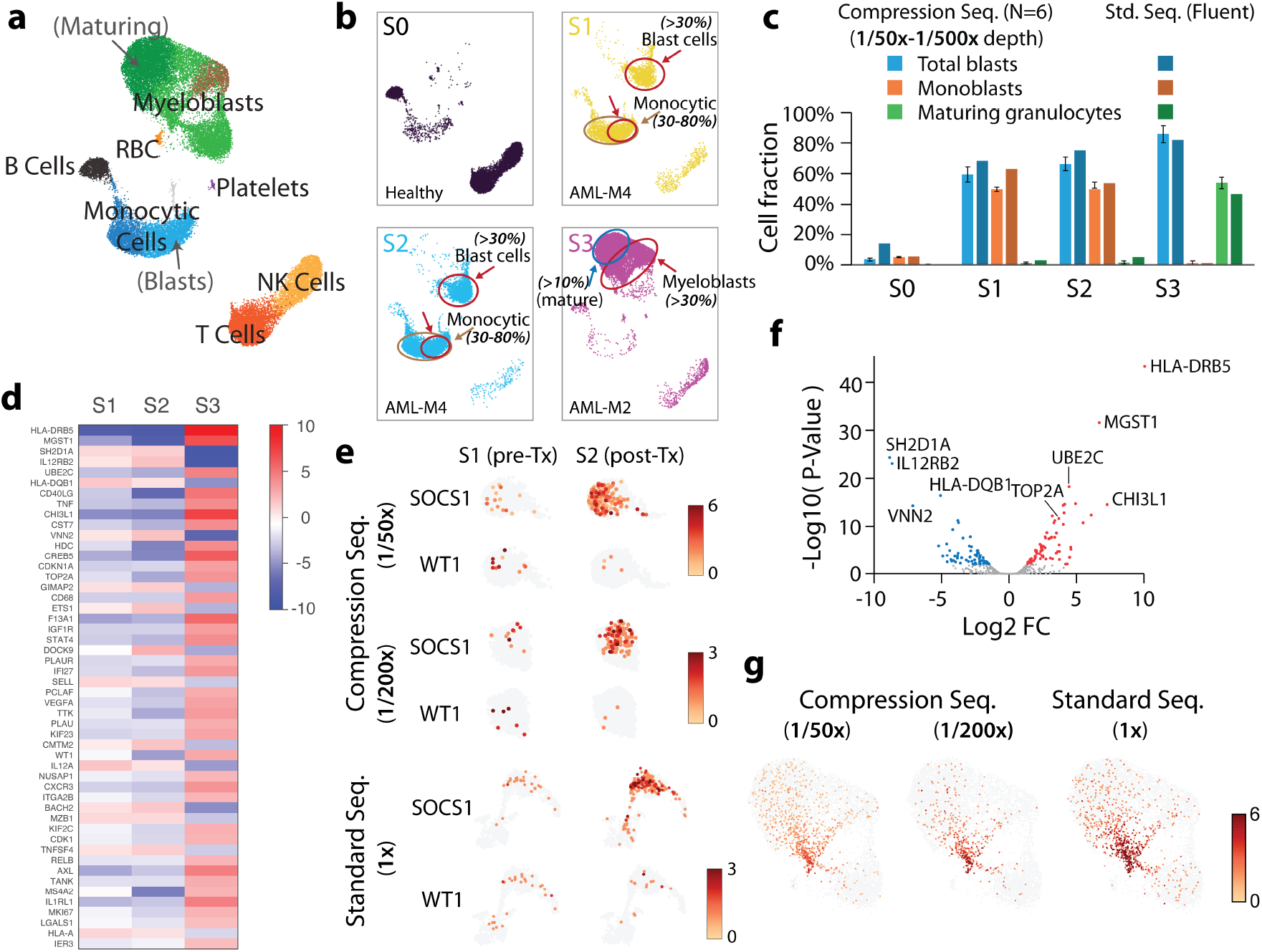
Compression Sequencing enables affordable AML diagnostics with in-depth transcriptomic analysis. **(a)** Aggregated dataset of 4 PBMC scRNA-seq samples: 1 healthy (S0), 3 with AML diagnosis (S1-S3), prepared using Fluent (now Illumina) PIP-seq kits, followed by Compressed Shallow Sequencing (1/60x depth of standard assay, 600 reads/cell vs 38,000 reads/cell). See Extended Data Fig. 7a for sample details. Cell lineages are assigned using marker-based annotation; arrows indicate cell sub-populations. (**b**) UMAP for individual samples. Labels indicate cell sub-population and fractions, along with clinical criteria for AML-M2 (AML with maturation) and AML-M4 (AMML) subtype diagnosis by FAB criteria. (**c**) Bar graph showing cell fractions measured by Compression Sequencing, compared to standard Fluent assay. Error bars indicate standard deviation from N=6 measurements, at 1/50x, 1/60x, 1/100x, 1/200x, 1/300x and 1/500x sequencing depth. (**d-g**) In-depth transcriptomic analysis in disease cell sub-populations using Compression Sequencing. (**d**) Differential expression profiles (top 50 genes) in myeloblasts across three AML samples (S1-S3). (**e**) Examples of differential expression profiles in paired AML-M4 samples before (S1) and after (S2) treatment, comparing Compression Sequencing (at 1/50x and 1/200x depth) with standard assay. (**f**) Volcano plot of DEGs in AML myeloblasts (S3), showing upregulated genes in immune regulation, active cell cycling, or associated with poor prognosis. (**g**) Identification of cell sub-population showing active cell cycling program in AML-M2 sample (S3). See Methods for analysis details.

We performed Compression Sequencing on the Fluent scRNA-seq libraries, using a 1,000-gene immunology marker panel (952 genes, 1084 total targets), which allows capture of AML-related genes for accurate cell type assignment and downstream gene expression profiling. Compressed Shallow Sequencing (600-800 reads/cell, 1/50-60x) showed consistent three-compartment cell clustering profile between two technical replicates and in high agreement with Fluent dataset (ARI=080-0.82, Fig. 5a, Extended Data Fig. 7, Supplementary Table 5). Further sub-sampled datasets showed similarly accurate cell clustering in mature cell types (ARI=0.72-0.83, accuracy 85-92%) down to a very shallow depth of 80 reads/cell (1/500x reduction from Fluent dataset, Extended Data Fig. 7d-e, Supplementary Table 5).

We next asked if Compressed Shallow Sequencing can accurately identify AML blasts and report disease subtypes. The 1,000-gene marker panel allowed transcriptomic assignment of monoblasts and maturing myeloblasts using multiple marker genes (Fig. 5a). Following the FAB revised criteria^55,56^: >30% total blasts, 30-80% monocytic cells for M4 subtype, <20% monocytic cells and >10% maturing granulocytes for M2 subtype (all fractions in non-erythroid cells), our results correctly reproduced AML subtypes in all three samples (Fig. 5b). Additionally, we observed stable cell fractions (total blasts, monocytic cells, maturing myeloblasts) from sub-sampled datasets down to 1/500x reduced sequencing depth (Fig. 5c), showing promise for accurate and affordable single-cell AML diagnostics.

In addition to cell fraction profiling, Compressed Shallow Sequencing (1/200x depth) allows comprehensive transcription program analysis in disease cell populations (Fig. 5d). For example, differential analysis between paired pre- and post-treatment M4 samples (Fig. 5e) showed significant upregulation of SOCS1 in the post-treatment blasts with no evidence of blast clearance, suggesting a therapy-induced transcriptional modulation. Additionally, slightly lower expression of MRD marker WT1 in the post-treatment population suggests effective treatment. In the M2 sample, we observed elevated levels of HLA-DRB5, suggesting potential immune regulation; as well as several upregulated genes associated with higher stemness and poor prognosis, e.g. CHI3L1, MGST1, UBE2C (Fig. 5f). Finally, we identified an actively cycling myeloid sub-population, characterized by high expression of cell-cycle genes (MKI67, TOP2A, CDK1, UBE2C, TK1), suggesting proliferative pressure and higher risk of residual disease persistence (Fig. 5g).

In summary, Compression Sequencing even at very shallow depth (1/200-500x of standard assay) preserves accurate cell state identification, correct disease subtype classification in AML disease samples, allowing accurate scRNA-seq disease testing. Beyond standard diagnostics, our method further allows in-depth transcriptomic profiling in disease cell sub-populations, providing valuable insights that can guide clinical treatment and monitoring strategies. Notably, despite very shallow sequencing depth, our method shows similar sensitivity as the reference dataset (Fig. 5e,g), and allows quantitative gene profiling across >1,000-fold of average abundance or differential expression. Although demonstrated in PBMC samples, our method is applicable to bone marrow or other sample types. Finally, we estimate the sequencing cost to be <$5 per sample (20,000-30,000 cells), enabling affordable single-cell diagnostics.

## Discussion

In this work, we reported a new, information science-inspired method, Compression Sequencing, that performs dynamic range reduction in complex nucleic acid libraires, and allows ultra-sensitive and scalable sequencing. Our method builds on Compression PCR (cPCR), a “molecular information encoder” that performs a logarithmic transform of nucleic acid sequence abundances, with a high accuracy, wide dynamic range (5+ logs), and in an unbiased fashion across hundreds to thousands of genes. By eliminating highly redundant reads, our method smartly redistributes sequencing budget to rare genes and transcripts, and achieves 100x higher sequencing efficiency in scRNA-seq samples. This 100x improvement translates to: (i) ultra-sensitive “Deep Sequencing”, revealing more transcripts and DEG detection in rare genes; and (ii) cost-effective “Shallow Sequencing”, profiling scRNA-seq samples at 200-500x lower cost (i.e. or similarly higher scale), while maintaining gene sensitivity, cell state accuracy, and disease subtype diagnosis.

Compression Sequencing enables high-quality, massively scalable single-cell assays for comprehensive functional genomics, drug x genetic screens, or building generalizable virtual cell models. For example, a whole-genome perturb-seq experiment (20,000 genes, 500 cells each) would require 10 M cells and an estimated >$1 M total cost. Current methods had to strike a touch balance between sensitivity, coverage and affordability; or resort to phenotypic enrichment based on a small number of selected markers. Building predictive and generalizable cellular foundation models takes even larger, high-quality training datasets (30-100 M cells), and are particularly sensitive to gene loss and batch effects. By improving sequencing efficiency and detection sensitivity over rare transcripts, Compression Sequencing can greatly reduce sequencing cost and potentially enable single-batch at massive scale (10-100 M cells) required for such studies (e.g. profiling 10 M cells at <$10,000 total cost).

Alternatively, using Deep Sequencing combined with microplate-based single-cell capture approaches (e.g. smart-seq3xpress^57^) allows in-depth analysis of rare gene expression patterns with true single-cell resolution, that requires >1 M reads/cell using standard approach^21^, and remains inaccessible at a large scale. Notably, the degree of compression can be fine-tuned by controlling limiting primer concentration and cPCR reaction conditions, allowing an adjustable balance between sensitivity, cell throughput and sequencing cost reduction.

Compression Sequencing shows the promise of affordable scRNA-seq disease testing in the clinic. With 200x reduced sequencing depth, our results reproduced AML clinical subtypes, and in addition allowed transcriptomic profiling over a 1,000-gene panel. This translates to $<5 sequencing cost (for 20,000-30,000 cells per sample) and $<10 estimated total cost per patient, allowing affordable disease diagnostics with comprehensive transcriptomic profiles.

High dynamic range and associated high sequencing cost pose a common challenge in many genomic and biomedical assays beyond scRNA-seq. Compression Sequencing transforms molecular abundances in complex libraries and is widely applicable to any of such systems, e.g. spatial transcriptomics^58,59^, viral diagnostics^60^ or plasma proteomics^61–63^. Future development of our method will extend to whole-transcriptome coverage, and integrate with probe-based single-cell libraries, spatial transcriptomics^58,64,65^ and multi-omic profiling^66–69^. We envision that Compression Sequencing will enable broad applications from basic research to clinical studies, including large-scale functional genomics, drug discovery, organism and development cell atlas, AI virtual cell and disease modeling, as well as affordable single-cell diagnostics.

* We note that Compression Sequencing is a distinct method from compressed Perturb-seq, which measures multiple random perturbations per cell. Here, Compression Sequencing improves sequencing efficiency by transforming molecular abundance in complex libraries and is broadly applicable to many sequencing-based assays (including Perturb-seq).

